# De novo design of a peptide ligand for specific affinity purification of human complement C1q

**DOI:** 10.64898/2026.03.30.714096

**Authors:** Ryo Tsuchihashi, Misaki Kinoshita, Hiroaki Aino

**Author notes:** **<u>Correspondence</u>** Hiroaki Aino, **<u>Address</u>** ^a^ Kobe KIMEC Center Building, Minatojima-minamimachi 1-5-2, Kobe Chuo-ku, Hyogo, 650-0047, Japan. (These authors contributed equally to this work.) (Ryo Tsuchihashi, Misaki Kinoshita).

## Abstract

Affinity purification is a essential technique for isolating highly purified proteins; however, generating affinity ligands require significant time and financial investment. To address these limitations, this study proposes a novel affinity chromatography method utilizing *in silico*-designed cyclic peptides as ligands. Targeting Complement C1q (C1q), a plasma protein that plays crucial roles in classical complement pathway, we employed the biomolecular structure prediction model, AlphaFold2, to design specific binding cyclic peptides. Based on these designs, we synthesized lariat-type cyclic peptides characterized by disulfide cyclization and biotinylation, which were subsequently immobilized on streptavidin carriers. Performance tests confirmed that the resulting column specifically captured C1q, allowing for elution via a standard NaCl concentration gradient. Notably, high selectivity was preserved even in the presence of plasma, underscoring the ligand’s practical robustness. By overcoming traditional constraints through (1) rapid and simple design, (2) high specificity, and (3) universal versatility without genetic modification, this *de novo* design strategy represents a potential breakthrough in protein purification technologies.

**Highlights:**

- AI-driven de novo design generated a specific cyclic peptide ligand for Complement C1q
- The synthetic ligand enabled one-step purification of Complement C1q directly from human plasma
- Mild elution conditions preserved the target’s oligomeric structure and native interactome
- This label-free strategy offers a rapid, low-cost alternative to antibody-based chromatography

## Context

## 1. Introduction

The isolation of high-fidelity proteins is a cornerstone of biotechnology, underpinning advancements from fundamental biology to pharmaceutical development. While affinity chromatography represents the gold standard for this purpose [1], the development and deployment of specific ligands remain significant bottlenecks. Conventional antibody generation relies on laborious and time-consuming biological screening processes, such as in vivo immunization or phage display [2,3]. Furthermore, while epitope tagging (e.g., His or FLAG tags) offers convenience, it necessitates genetic modification, precluding the purification of endogenous proteins from native tissue or clinical samples [4,5]. Consequently, a universal, label-free strategy capable of rapidly generating low-cost ligands for any given target remains a critical unmet need in protein science.

In this study, we address this challenge using the purification of Complement C1q (C1q) from human plasma as a model system. C1q, a pivotal bridge between innate and adaptive immunity, is strongly implicated in autoimmune disorders like systemic lupus erythematosus (SLE) and neurodegenerative diseases [6,7]. C1q is a structurally complicated, 460 kDa heterooligomeric complex composed of 18 polypeptides [8]. Conventional physicochemical fractionation is inefficient and multi-staged, resulting in poor yields, while existing antibody columns are cost-prohibitive and often require acidic elution that compromises the structural integrity of such labile complexes [9,10,11].

We leverage the paradigm shift in protein structure prediction by AlphaFold2 [12] to revolutionize affinity purification. While in silico de novo design algorithms have primarily been explored in the context of therapeutic inhibitors, we repurposed this technology for “purification ligand engineering”. This method enables more efficient and cost-effective protein purification than traditional approaches. Guided by C1q crystal structure data [13], we employed in silico directed evolution to design and chemically synthesize C1q-specific cyclic peptides. When immobilized on a chromatographic resin, these synthetic ligands successfully captured C1q directly from the complex matrix of human plasma. Crucially, the target was eluted under mild conditions that preserved its physiological activity and oligomeric structure. This study demonstrates that AI-driven molecular design can provide a rapid, universal, and cost-effective affinity purification platform applicable to endogenous proteins, overcoming long-standing barriers in the field.

## 2. Materials and methods

### 2.1. Materials

C1q was purified from Cohn’s fractionated human blood plasma by multi-step ion-exchange chromatography procedure. Unless otherwise specified, analytical grade reagents were purchased from FUJIFILM Wako Pure Chemical Corporation (Osaka, Japan) or Sigma-Aldrich (St. Louis, MO, USA). The ACQUITY UPLC BEH 200 SEC column and BEH 200 SEC Protein Standard Mix were purchased from Waters Corporation (Milford, MA, USA). NuPAGE 4–12% Bis-Tris gels, PVDF membranes, nitrocellulose membranes, LDS Sample Buffer, and Reducing Agent were purchased from Thermo Fisher Scientific (Waltham, MA, USA). HBS-EP+ running buffer, Sensor Chip SA, Amersham ECL Prime, and HiTrap Streptavidin HP columns were purchased from Cytiva (Marlborough, MA, USA). Standard human plasma was purchased from Sysmex (Kobe, Japan). InstantBlue Coomassie protein stain was purchased from Abcam (Cambridge, UK), and Bullet Blocking One was purchased from Nacalai Tesque (Kyoto, Japan). Primary anti-C1qC antibody was purchased from Abcam, and HRP-conjugated secondary antibody was purchased from Cytiva. The custom-designed cyclic peptide was synthesized by Eurofins Genomics (Tokyo, Japan).

### 2.2. In silico design of C1q affinity cyclic peptides

The C1q-affinity peptides were designed using an in-house AlphaFold2-based design pipeline, specifically optimized for the conformational prediction of cyclic peptides. Computational simulations were accelerated using NVIDIA T4 GPUs. Following an optimization trajectory, we identified a lead candidate (designated as Rank 1) with the primary sequence DPYGDYNPKYYPE. The structural confidence of the predicted C1q-peptide complex was evaluated using the predicted Local Distance Difference Test (pLDDT), which yielded a robust score of 85.28, indicating a high degree of local model reliability.

### 2.3. SPR measurements

The binding kinetics between C1q and the synthetic peptides were measured using a Biacore X100 system (Cytiva) at 25°C. All experiments were conducted in HBS-EP+ as the running buffer. Biotinylated peptides of 0.4 μg/mL were immobilized onto a Sensor Chip SA to achieve a target density of 300 resonance units. For the binding assays, C1q was prepared in a concentration series ranging from 1 ng/mL to 10 μg/mL. The analyte was introduced using a single-cycle kinetics approach with a contact time of 60 seconds for each concentration to obtain the representative sensorgrams.

### 2.4. Affinity column chromatography

The C1q-binding peptide column was prepared by immobilizing a biotinylated synthetic peptide onto a streptavidin-coupled solid phase. Briefly, 5.7 mg of the peptide was dissolved in 1 mL of Milli-Q water, from which a 0.5 mL aliquot was further diluted with 0.6 mL of PBS. The final peptide concentration was verified by spectrophotometric analysis using a spectrophotometer. A streptavidin column was equilibrated with at least 10 column volumes (CV) of PBS, followed by the application of 1.1 mL of the prepared peptide solution via a syringe. The column was incubated overnight at room temperature to ensure maximal biotin-streptavidin binding, while a control column was similarly prepared by applying PBS in place of the peptide. After incubation, the immobilization efficiency was estimated by measuring the absorbance of the flow-through fraction collected during a 1 mL PBS wash. The resulting C1q-binding peptide column was washed with 10 mL of PBS and stored at 4 °C until further use.

Chromatographic separation was conducted using an ÄKTA pure 150 system. The column with or without peptide ligand immobilization was equilibrated with diluted PBS as the equilibration phase, and samples were subsequently injected. Then, a linear gradient elution over 5 column volume (CV) was applied with fixed volume fractionation. To evaluate the binding capability of the peptide to purified C1q, C1q sample solution was diluted to 100 μg/mL using Milli-Q water in order to decrease the ionic strength, and eluted using a gradient from 12.5% to 100% Buffer B (1× PBS) against Buffer A (0.25× PBS). For experiments to examine the purification performance of peptide, standard human plasma was reconstituted in 1 mL of Milli-Q water, further diluted with an additional 5 mL of Milli-Q water, and supplemented with 100 µL of C1q. These C1q-spiked plasma samples were eluted with a linear gradient from 0% to 100% Buffer B (1× PBS) against Buffer A (0.33× PBS) with fixed volume fractionation.

### 2.5. SDS-PAGE, Immunoblotting, and Dot Blot Analysis

Protein purity and subunit composition were assessed by SDS-PAGE using NuPAGE 4–12% Bis-Tris gels. Samples were prepared in LDS Sample Buffer with a Reducing Agent and heat-denatured prior to electrophoresis at 70 °C. Proteins were visualized by Instant Blue staining or electro-transferred onto PVDF membranes using the iBlot 2 Dry Blotting System. For dot blot analysis, 1 μL of each sample was spotted onto a nitrocellulose membrane and air-dried. Both PVDF and nitrocellulose membranes were blocked with Bullet Blocking One for 5 min. The membranes were then incubated with a primary anti-C1qC antibody, followed by an HRP-conjugated secondary antibody. Chemiluminescent signals were developed using Amersham ECL Prime and captured with an Amersham Imager 600. Signal intensities for the dot blots were quantified using the Protein Array Analyzer plugin for ImageJ software [14].

### 2.6. UPLC-SEC analysis of purified C1q bound to peptide column

The oligomeric profile of C1q sample aliquot was characterized using an ACQUITY UPLC system equipped with a BEH 200 SEC column. Following clarification through a 0.22 μm spin filter, a 2 μL aliquot was injected and eluted isocratically at a flow rate of 0.3 mL/min using a mobile phase consisting of 50 mM Tris-HCl (pH 7.4), 150 mM NaCl, and 5 mM CaCl_**2**_. The column and sample manager were maintained at 30 °C and 10 °C, respectively. Protein elution was monitored by UV absorbance at 280 nm and 220 nm, with molecular weights estimated using a BEH 200 SEC Protein Standard Mix.

## 3. Results

### 3.1. In Silico Design of C1q-Affinity Cyclic Peptides

To engineer high-affinity ligands targeting C1q, we employed the evolutionary optimization algorithm using AlphaFold2 for structural prediction. The crystal structure of the recombinant globular heads of C1q (gC1q), comprising chains A, B, and C (PDB ID: 5HKJ) [13], served as the structural reference. Using the amino acid sequences derived from this template as a reference structure, we generated a library of candidate binders. From the output, the sequence “DPYGDYNPKYYPE” exhibited the highest predicted ranking score and was selected as the lead candidate (Fig. 1).

**Figure. 1.**
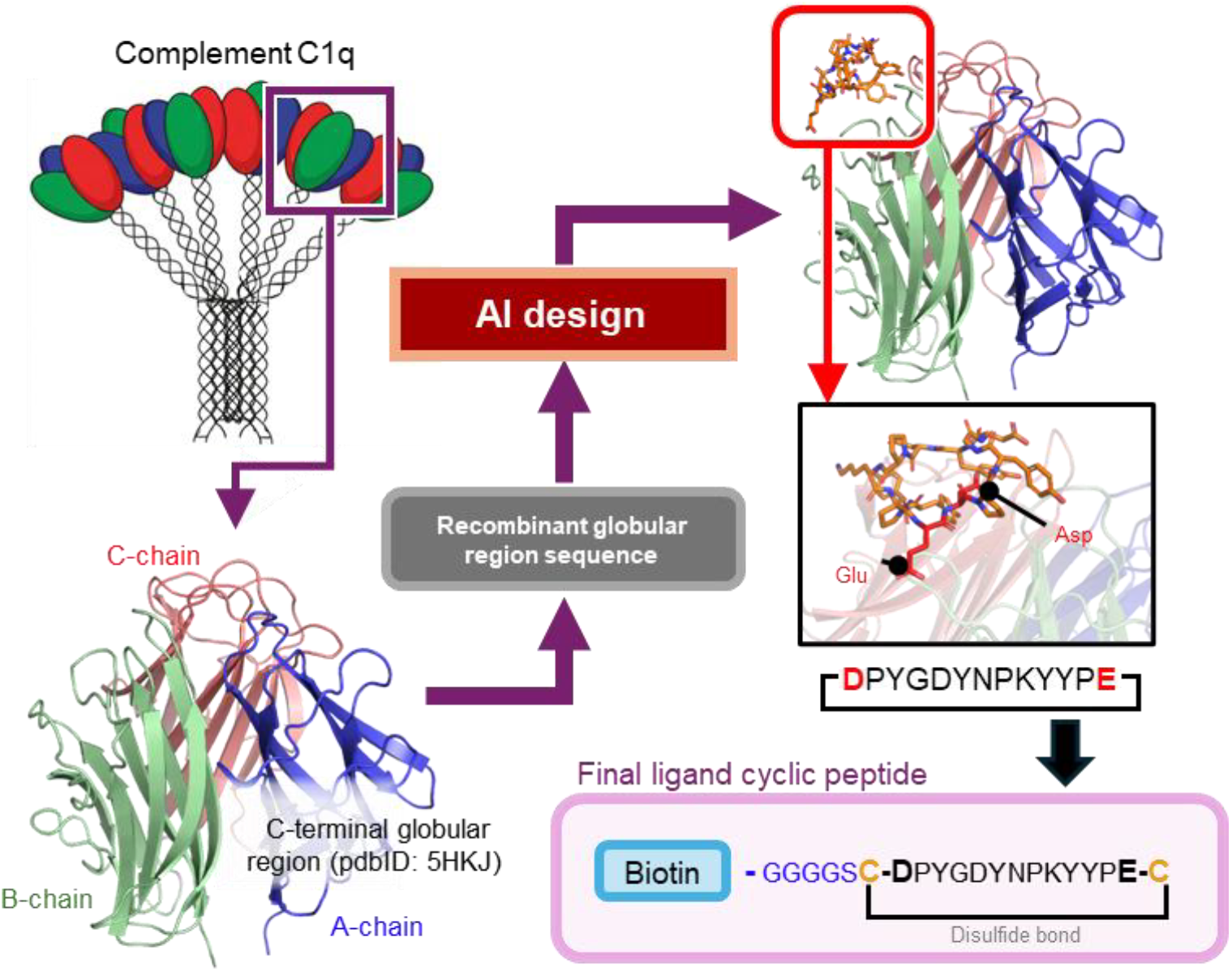
Design scheme of C1q affinity cyclic peptide. C1q is a large 18-chain oligomer, in which C1q A, B, and C chains constrain trimer, and six such trimers constitute the full C1q native structure. To design the C1q-affinity cyclic peptide, the crystal structure of the recombinant C1q globular domain (gC1q) was utilized as the target (PDB ID: 5HKJ), as indicated by the purple square in the schematic illustration. The sequence of this recombinant gC1q served as the reference sequence for initial AlphaFold2 reference structure, and the sequences around N- and C-termini were excluded to ensure the peptide-binding site remained broadly solvent-exposed. For surface immobilization in the SPR measurements and affinity chromatography, designed cyclic peptide with highest score is linearized by cleaving the peptide bond between D1 and E13. Cysteine residues were then appended to both termini to allow for re-cyclization via a disulfide bond. Finally, a biotin tag with a Gly-Ser linker (GGGGS) was conjugated to the N-terminal cysteine.

The initially designed scaffold featured a fully closed, head-to-tail cyclic backbone, which lacks a functional handle for resin immobilization. So, we linearized the backbone at the D1–E13 junction and re-engineered the peptide to maintain its cyclic constraint via a disulfide bond formed between newly introduced terminal cysteine residues. To facilitate immobilization, a biotin functional moiety was conjugated to the N-terminus via a glycine-serine (GS) linker (Fig. 1).

### 3.2. Evaluation of C1q Binding Affinity and Chromatographic Performance

To validate the interaction between the synthetic peptide and native C1q, we performed Surface Plasmon Resonance (SPR) analysis. Sensorgrams demonstrated distinct, concentration-dependent binding responses of C1q to the peptide-immobilized chip (Fig. 2A). Specifically, upon immobilizing approximately 300 Response Units (RU) of the peptide, the injection of 10 ug/mL C1q yielded a binding response of approximately 1000 RU. Notably, the dissociation phase exhibited rapid signal decay, indicating a relatively moderate binding affinity. We attribute this kinetic profile to minor structural deviations from the in silico energetic optimum, likely induced by the re-engineering of the cyclic constraint and the addition of the linker-biotin moiety. Nevertheless, the results confirm that the core geometry remains sufficiently intact to recognize the target C1q structure, despite the observed reduction in affinity. Given the confirmed binding activity, we next assessed the utility of the designed peptide as an affinity ligand for chromatographic purification using a streptavidin-conjugated column. Since preliminary SPR data suggested a salt-sensitive binding mechanism, we optimized the loading buffer to low-ionic strength conditions and established an elution protocol using a linear gradient increasing to 1xPBS. In control experiments using a peptide-free streptavidin column, C1q failed to bind and was recovered exclusively in the flow-through fraction (Fig. 2B, gray solid line). In contrast, the peptide-functionalized column exhibited significant retention of C1q, which eluted as a distinct peak at a conductivity of 8 mS/cm (Fig. 2B, blue solid line). This chromatographic behavior aligns with the SPR kinetics, demonstrating that the immobilized peptide specifically captures C1q and allows for elution via ionic strength modulation. With a total peptide load of approximately 2.85 mg, the column capacity was in significant molar excess relative to the applied C1q sample, ensuring the efficient binding to the input C1q.

**Figure. 2.**
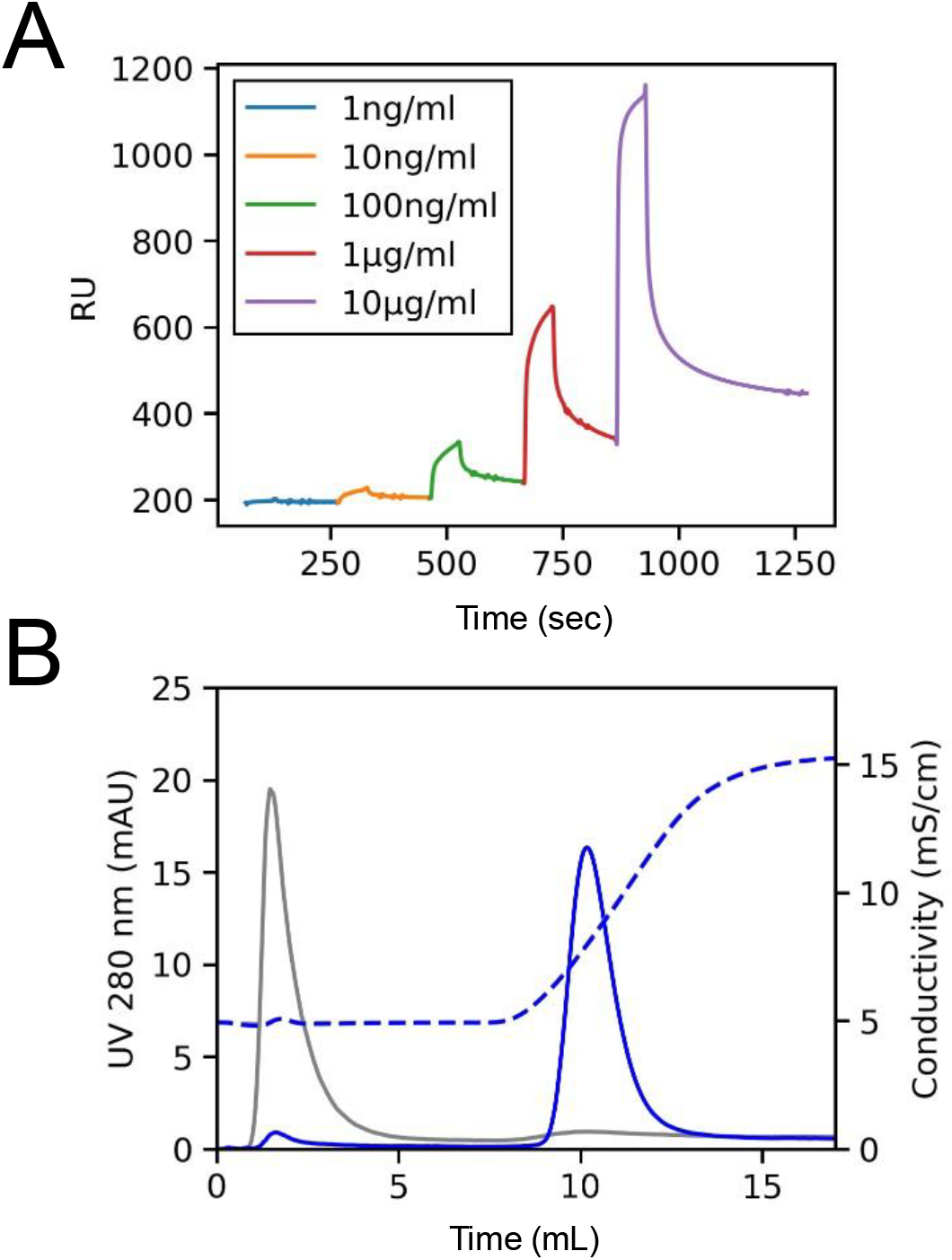
Affinity of the designed cyclic peptide for native C1q. (A) SPR sensorgrams for the interaction between designed cyclic peptide and C1q. Biotinylated cyclic peptide used as ligand was immobilized on the streptavidin coated sensor chip, and native C1q purified from human plasma at desired concentrations was injected. (B) Evaluation of interaction between cyclic peptide and C1q using affinity chromatography. Purified C1q was applied to a streptavidin column either without (gray) or with (blue) the immobilized biotinylated cyclic peptide. Solid and dashed lines represent UV absorbance at 280 nm and conductivity, respectively.

### 3.3. Selective Purification of C1q from Complex Biological Matrix

Following validation with purified standards, we investigated the capability of the cyclic peptide-functionalized column to selectively isolate C1q from a complex biological milieu. Standard human plasma spiked with C1q was applied to the column, yielding a distinct elution peak at a conductivity of approximately 8 mS/cm, which closely resembled the profile of the purified standard (Fig. 3A). To verify the presence of the eluted species, fractions were first analyzed via Dot Blot using an anti-C1q C-chain antibody. Intense specific signals were detected exclusively in the elution fraction, with negligible antigen in the flow-through, confirming the successful capture of C1q (Fig. 3B). Subsequently, the components of the elution fraction were characterized by SDS-PAGE (Coomassie Brilliant Blue staining) and Western blotting (Fig. 3C). While the flow-through contained the bulk of the plasma proteome, the eluate displayed a distinct electrophoretic signature. Under reducing conditions, C1q typically resolves into two characteristic bands: a higher molecular weight band corresponding to the co-migrating A chain (∼34 kDa) and B chain (∼32 kDa), and a lower molecular weight band corresponding to the C chain (∼27 kDa). The doublet observed in the 20 to 38 kDa range in our eluate is consistent with this signature (Fig. 3C white and black triangles, respectively), and immunoblotting using an anti-C1q C-chain antibody confirmed this identity.

**Figure. 3.**
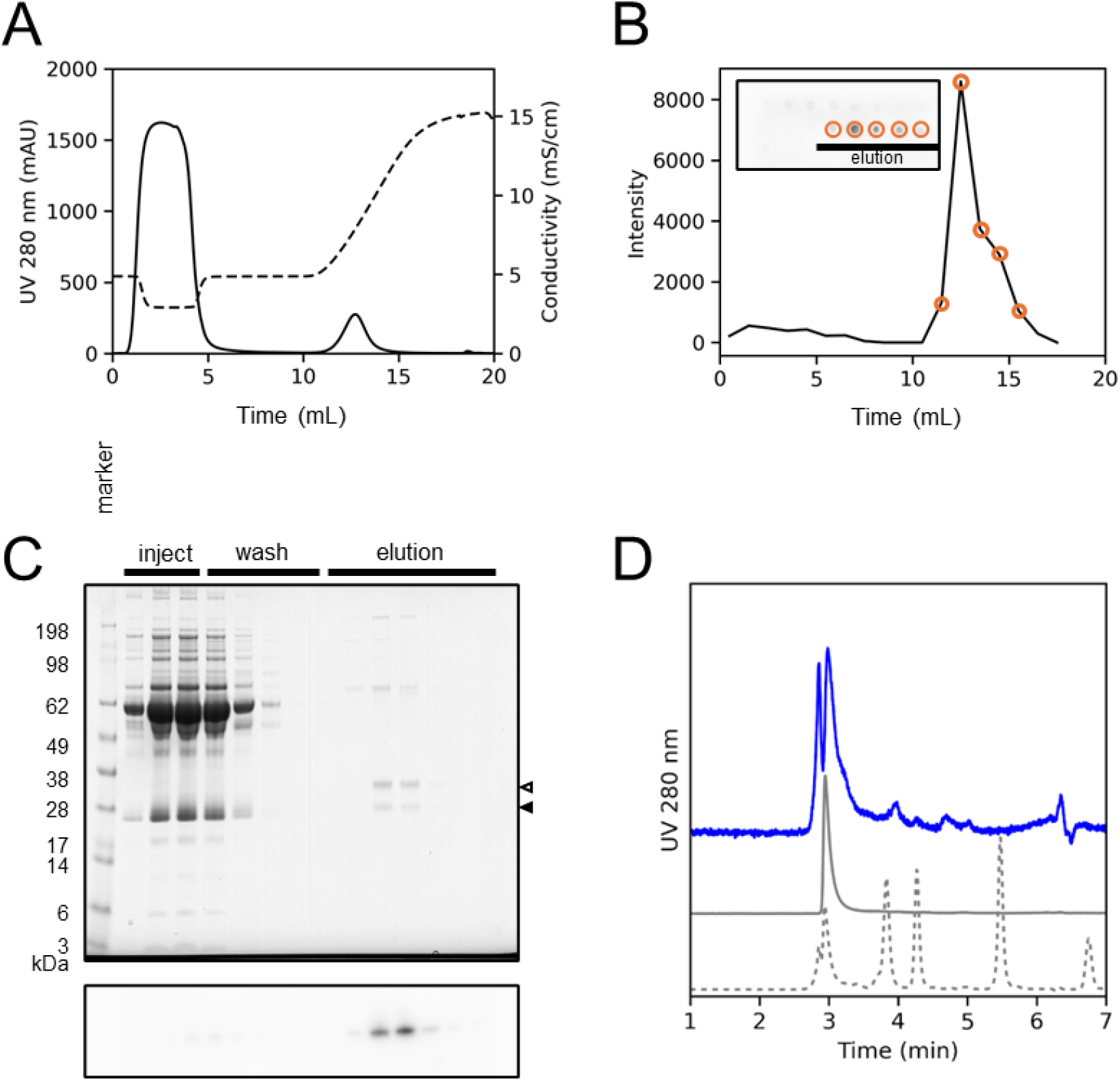
Specific affinity of designed cyclic peptide for C1q. (A) Affinity chromatography using a cyclic peptide-immobilized column with C1q-spiked human plasma. Solid and dotted lines represent UV absorbance at 280 nm and conductivity, respectively. (B) Dot blot analysis for the detection of C1q protein across fractions. Signal intensities are plotted against elution volume of the chromatogram represented in (A), with the raw chemiluminescence image shown in the inset. (C) Analysis of elution fractions by SDS-PAGE and Western blotting. The upper panel shows the CBB-stained gel, and the lower panel shows the Western blot using an anti-C1q C-chain antibody. The C1q C-chain was specifically detected in the elution fractions (black triangle represents the C1q C-chain, and white triangle represents both C1q A- and B-chains), where minimal protein impurities were observed in the CBB stained gel. (D) UPLC-SEC chromatograms of eluted peptide bound fraction (solid blue line), reference purified C1q protein (solid gray line), and molecular weight standard (dotted line). The standards eluted in decreasing order of molecular weight (increasing retention time) as follows: thyroglobulin (∼669 kDa), IgG (∼150 kDa), BSA (∼66 kDa), myoglobin (∼17 kDa), and uracil (small molecule marker).

Notably, several additional bands exceeding 62 kDa were observed in the elution fraction (Fig.3C). Given their co-elution behavior with the C1q subunits, these likely represent known C1q-associated proteins—such as fibronectin (∼220 kDa) [15] and vitronectin (∼75 kDa) [16]—that form stable complexes with C1q in circulation. Further UPLC analysis of the elution fraction (Fig. 3D) also revealed additional peaks alongside the primary C1q peak. Specifically, the standard C1q (solid gray line) eluted as a single peak between thyroglobulin (∼669 kDa) and IgG (∼150 kDa), and a corresponding prominent peak was clearly confirmed in the elution fraction. Furthermore, the elution profile showed one peak with a larger molecular size than the standard C1q, along with several other peaks, indicating the co-existence of proteins with lower molecular weights. This implicated that these species were off-target proteins co-purified through their interaction with the captured C1q; however, the possibility that these proteins were recovered due to non-specific binding to the column matrix or the peptide ligand cannot be ruled out at this stage.

## 4. Conclusion

In this study, we successfully leveraged AI-driven in silico de novo design to engineer a cyclic peptide ligand capable of specifically capturing the native C1q heterooligomer. The designed ligand enabled the one-step purification of C1q from plasma—a complex matrix teeming with contaminants— under mild conditions that maintained the target’s structural integrity. This methodology effectively circumvents the intrinsic limitations of traditional affinity purification, specifically the substantial time and costs associated with antibody development.

Nevertheless, challenges regarding the universal application of this technology remain. The success of the current design was partially contingent on the absence of post-translational modifications (PTMs), such as glycosylation or phosphorylation, at the selected binding interface [17,18]. Since many secreted proteins undergo extensive glycosylation in endoplasmic reticulum and golgi apparatus [19], which can sterically hinder designed interactions, future strategies must integrate structural data with PTM databases to proactively identify and avoid modified residues. Furthermore, the potential for binding interfaces to be occluded by endogenous partners in biological fluids must be considered. While the current process still entails a degree of trial and error, we anticipate that rapid advancements in AI algorithms will soon overcome these structural and environmental constraints [20].

Of particular significance is the observation that known C1q-interacting proteins were co-eluted during the purification process. This behavior suggests that the ligand captures the target as a non-covalent complex, preserving its physiological associations. Consequently, this peptide-based strategy transcends simple protein isolation; it serves as a robust platform for interactome analysis that reflects the native biological state.

In conclusion, the generative AI-driven molecular design technologies that are currently revolutionizing drug discovery are poised to catalyze a parallel paradigm shift in the foundational biotechnology of protein purification. We envision this platform as a cornerstone of next-generation protein science, offering a “rapid, cost-effective, and high-purity” solution applicable to the entire proteome.

## CRediT authorship contribution statement

**Ryo Tsuchihashi:** Conceptualization, Methodology, Formal analysis, Investigation, Visualization. **Misaki Kinoshita:** Conceptualization, Investigation, Writing – original draft, Visualization. **Hiroaki Aino:** Supervision, Writing – review & editing.

